# Supervised fine-tuning of pre-trained antibody language models improves antigen specificity prediction

**DOI:** 10.1101/2024.05.13.593807

**Authors:** Meng Wang, Jonathan Patsenker, Henry Li, Yuval Kluger, Steven H. Kleinstein

## Abstract

Antibodies play a crucial role in adaptive immune responses by determining B cell specificity to antigens and focusing immune function on target pathogens. Accurate prediction of antibody-antigen specificity directly from antibody sequencing data would be a great aid in understanding immune responses, guiding vaccine design, and developing antibody-based therapeutics. In this study, we present a method of supervised fine-tuning for antibody language models, which improves on previous results in binding specificity prediction to SARS-CoV-2 spike protein and influenza hemagglutinin. We perform supervised fine-tuning on four pre-trained antibody language models to predict specificity to these antigens and demonstrate that fine-tuned language model classifiers exhibit enhanced predictive accuracy compared to classifiers trained on pretrained model embeddings. The change of model attention activations after supervised fine-tuning suggested that this performance was driven by an increased model focus on the complementarity determining regions (CDRs). Application of the supervised fine-tuned models to BCR repertoire data demonstrated that these models could recognize the specific responses elicited by influenza and SARS-CoV-2 vaccination. Overall, our study highlights the benefits of supervised fine-tuning on pre-trained antibody language models as a mechanism to improve antigen specificity prediction.

**Author Summary:** Antibodies are vigilant sentinels of our adaptive immune system that recognize and bind to targets on foreign pathogens, known as antigens. This interaction between antibody and antigen is highly specific, akin to a fitting lock and key mechanism, to ensure each antibody precisely targets its intended antigen. Recent advancements in language modeling have led to the development of antibody language model to decode specificity information in the sequences of antibodies. We introduce a method based on supervised fine-tuning, which enhances the accuracy of antibody language models in predicting antibody-antigen interactions. By training these models on large datasets of antibody sequences, we can better predict which antibodies will bind to important antigens such as those found on the surface of viruses like SARS-CoV-2 and influenza. Moreover, our study demonstrates the potential of the models to “read” B cell repertoire data and predict ongoing responses, offering new insights into how our bodies respond to vaccination. These findings have significant implications for vaccine design, as accurate prediction of antibody specificity can guide the development of more effective vaccines.

## Introduction

Recent advancements in natural language processing (NLP) have catalyzed the development of antibody language models, specialized deep learning architectures trained on vast datasets of antibody sequences [1–8]. These models leverage techniques such as masked language modeling and attention mechanisms [9,10] to encode the complex sequence-structure-function relationships inherent in antibodies [11,12] and hold promise to improve our understanding of immune responses by enabling high-throughput prediction of antigen specificity. Moreover, they offer a powerful framework for analyzing and interpreting large-scale antibody repertoire sequencing data [8], shedding light on the molecular mechanisms underlying immune system function and dysfunction.

Transfer learning, a cornerstone of modern machine learning, has emerged as a powerful paradigm for leveraging knowledge from one domain to improve performance in another. In the context of language models, transfer learning involves pre-training a neural network on a large dataset in a source domain and then fine-tuning it on a smaller dataset in a target domain, where labeled data may be scarce [13,14]. This approach capitalizes on the transferability of learned representations across related tasks or domains, enabling models to capture generic features that are transferable while adapting to task-specific nuances during fine-tuning without requiring extensive computational resources or labeled data. In the realm of antibody language models, fine-tuning offers a promising avenue for enhancing predictive accuracy and generalization across diverse antigen-specificity prediction tasks [6,8].

In this study, we investigated the efficacy of supervised fine-tuning of pre-trained antibody language models in predicting binding specificity to two key antigens: the SARS-CoV-2 spike protein and influenza hemagglutinin. By fine-tuning pre-trained models on labeled data specific to these antigens, we aimed to enhance predictive accuracy and generalization across diverse antibody sequences. We further applied the fine-tuned models to BCR repertoire data for influenza and SARS-CoV-2 vaccination to investigate their ability to capture changes induced by ongoing immune responses.

## Results

### Fine-tuning antibody language models for specificity prediction

To investigate the effect of supervised fine-tuning on predicting BCR specificity, we fine-tuned the last three layers of four pre-trained antibody language models, including antiBERTy [1], antiBERTa2 [3], BALM-paired [6], and ft-ESM2 [6] (**Table 1**), for binary binding status classification for SARS-CoV-2 spike (S) protein and influenza hemagglutinin (HA) (**Figure 1**). For performance comparison, we also trained supervised SVM on pre-trained model embeddings on the same task and data.

**Table 1.** Parameters of publicly available pre-trained antibody language models.

| Model | Parameters | Hidden Size | Intermediate Size | Attention Heads | Layers |
| --- | --- | --- | --- | --- | --- |
| antiBERTy | 25.76M | 512 | 2048 | 8 | 8 |
| antiBERTa2 | 202.64M | 1024 | 4096 | 16 | 16 |
| BALM-paired | 303.92M | 1024 | 4096 | 16 | 24 |
| ft-ESM2 | 652.36M | 1280 | 5120 | 20 | 33 |

**Figure 1.**
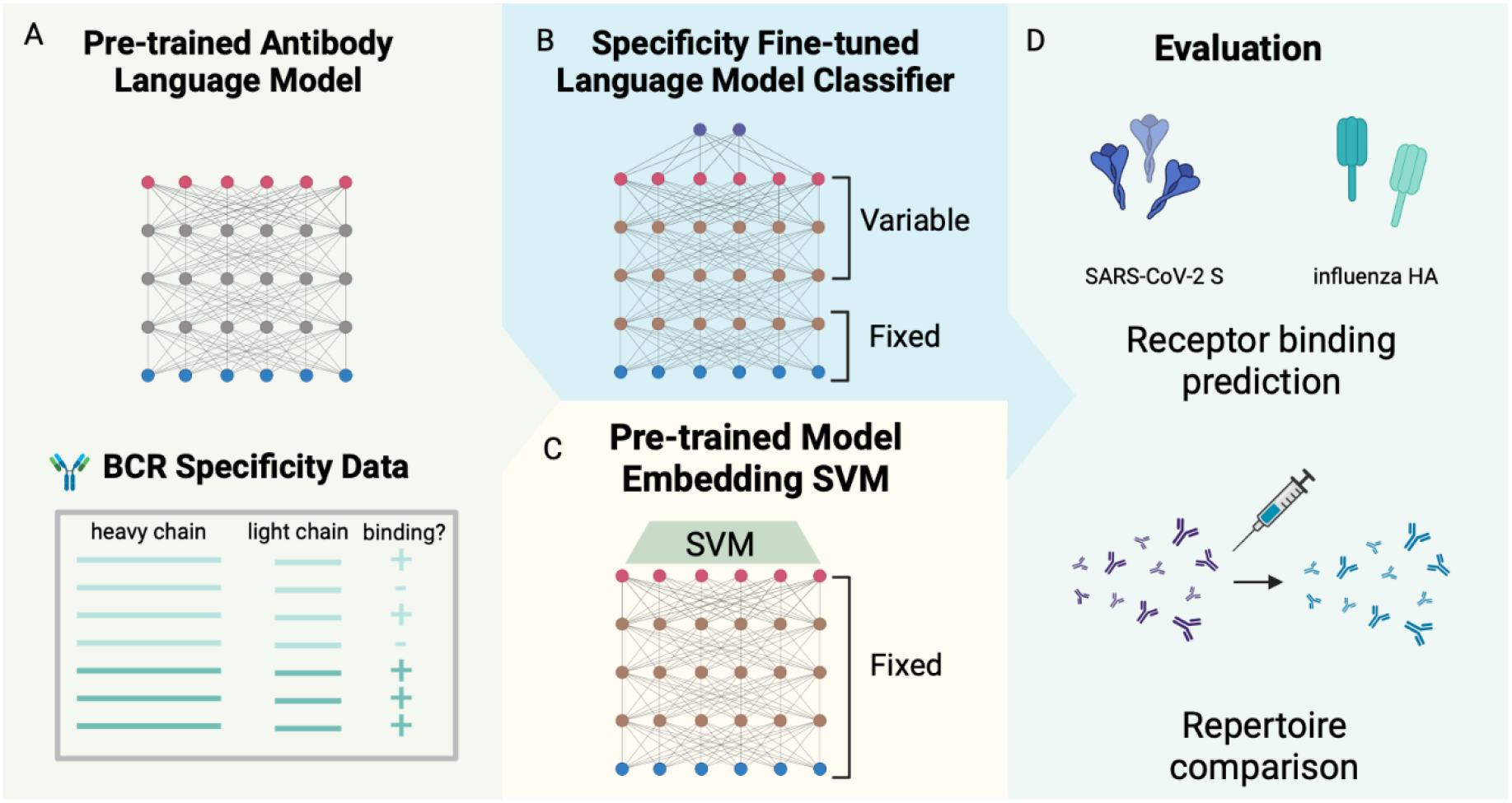
Fine-tuning antibody language model on receptor specificity prediction tasks. (A) Antibody language model-based specificity classifiers for SARS-CoV-2 S protein and influenza hemagglutinin were trained by (B) fine-tuning pre-trained antibody language models on specificity classification tasks, and (C) using supervised support vector machine classifiers on pre-trained model embeddings of BCR sequences. (D) The performance of the classifiers was evaluated on the cross-validation accuracy of receptor specificity prediction and comparison between longitudinal time points of repertoire datasets for SARS-CoV-2 and influenza vaccination.

Specificity of the models was evaluated on, in total, 15,539 and 5,514 paired full-length BCR sequences for S protein and HA fine-tuning classification task, respectively, from publicly available datasets with binding and donor/study labels (**Table 2**). To balance the classes, we sampled S protein non-binding sequences from pre-pandemic B cell repertoires, described in [15], and HA non-binding sequences from influenza vaccine non-responsive B cells [16]. The sampled non-binding sequences had similar V, J gene usage, CDR3 length, and somatic hypermutation frequency with the binding class (**Figure S1**).

**Table 2.** Data sources for antigen-specific antibody sequence and vaccination related BCR repertoire.

| Antigen | Data source | Type | Description | # Receptors |
| --- | --- | --- | --- | --- |
| Influenza HA | IEDB [21], downloaded on Dec 2023 | Sequence | Public database with curated BCR and epitope information, extracted human BCR sequences to influenza HA | 113 |
| Influenza HA | Turner 2020 [23]; McIntire 2024 (submitted) | Sequence | Human monoclonal antibodies sequences tested for binding to 2018 QIV HA | 1,551 |
| Influenza HA | Wang Y. 2023 [22] | Sequence | Curated database of human monoclonal antibodies to influenza HA | 1,311 |
| Influenza HA | Wang M. 2023 [16] | Sequence | Influenza vaccination non-responsive BCR sequences | 2,539 |
| Influenza HA | Wang M. 2023 [16] | Repertoire | Single-cell BCR sequencing on patients receiving seasonal influenza vaccination | 87,230 |
| SARS-CoV2 S | Wang M. 2024 [15] | Sequence | Collected public databases of human monoclonal antibodies to SARS-CoV-2 S protein | 15,539 |
| SARS-CoV2 S | Kim 2022 [18] | Repertoire | Single-cell BCR sequencing on patients receiving SARS-CoV-2 mRNA vaccine | 164,252 |

To evaluate the fine-tuned classifiers and the pre-trained model embedding classifiers, we used four-fold cross-validation (CV) with non-overlapping donors/studies between each train-test split and evaluated the performance of the models on the test split. Within the training split of each fold, we also performed hyperparameter selection for fine-tuning by further splitting a validation set (33%) from the train set or performing another three-fold cross validation to train the pre-trained embedding SVM.

### Supervised fine-tuning improves specificity prediction performance

As a performance baseline for specificity prediction, we evaluated the nested cross-validation performance of an SVM model on the embeddings from the four pre-trained antibody language models as well as the original ESM2 protein language model for SARS-CoV-2 S protein (**Figure 2A, Table S1**) and influenza HA (**Figure 2C, Table S2**) specificity prediction. We used different sequence inputs to generate the embeddings, including paired full length (FULL HL), full-length heavy chain (FULL H), paired CDR3 (CDR3 HL), and CDR3 heavy chain (CDR3 H). Consistent with previously reporting [15], the performance improved as we included longer sequences of the receptors for each antibody language model (FULL HL > FULL H > CDR3 HL > CDR3 H). For the full-length paired sequence input, ft-ESM2 performed the best across the language models for most of the evaluation metrics with an average CV test AUROC of 0.88 for S protein and 0.86 for HA.

**Figure 2.**
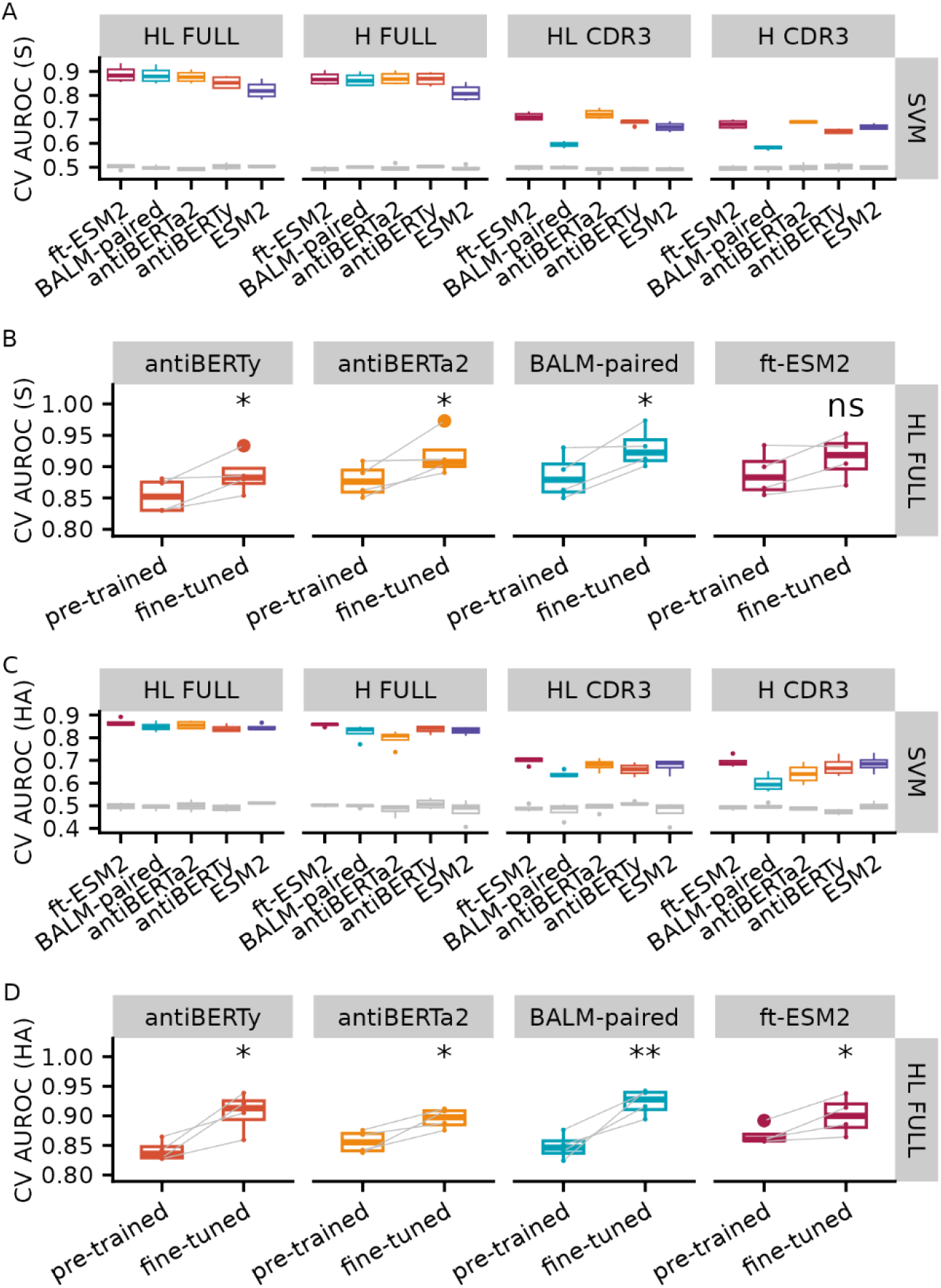
Specificity prediction performance for pre-trained model embedding SVM and fine-tuned antibody language models. Box plot of 4-fold cross validation AUROC for pretrained model embedding SVM in predicting binding to **(A)** SARS-CoV-2 S protein and **(C)** influenza HA with different sequence inputs. The gray box plots represent the random baseline by training the same model on shuffled labels. Comparison of CV AUROC between the pretrained embedding-based SVM model and fine-tuned language models on **(B)** SARS-CoV-2 S protein and **(D)** influenza HA binding data. Each line represents the test performance for one of the CV folds. Note that the pre-trained embedding based SVM model and fine-tuned models were trained and tested on the same data for each fold. Paired t-test was used to obtain the significance level of the increase in AUROC after fine-tuning (ns: p > 0.05, *: p <= 0.05, **: p <= 0.01).

We fine-tuned the four antibody language models by training the last three layers of the pre-trained model along with sequence classification head to predict the specificity of SARS-CoV-2 S protein (**Figure 2B, Table S3**) and influenza HA (**Figure 2D, Table S4**) using the full-length paired BCR sequences and evaluated the performance of fine-tuning on the test set by the same data split using the four-fold cross validation procedure as the pre-trained embedding procedure. For both antigens, we noticed an increase in the AUROC for all CV folds for fine-tuned classifiers compared with pre-trained embedding classifiers. We performed paired Wilcoxon-rank sum tests to examine whether the increases are significant and found that the increases are significant for all models except the ft-ESM2 after fine-tuning for S protein classification.

### Fine-tuning increases model attention at the CDR regions

Previous studies [2,6,17] have shown that protein language models trained on antibody sequences with the masked language model objective have increased self-attention activations on the locations of long-range structural contacts or functionally important regions for binding. To evaluate the effect of supervised specificity fine-tuning on the antibody language model self-attention activations [9], we randomly selected fifty antibodies specific for SARS-CoV-2 S protein and influenza HA from the training dataset, and computed the average intra-chain attention along the antibody heavy and light sequences across the last three layers of the four antibody language models before and after fine-tuning. We took the differences of the average intra-chain attention between the fine-tuned and pre-trained model and found an increase in average attention activations across all four models after fine-tuning in positions corresponding to the CDR regions, especially the CDR3 regions (**Figure 3, Figure S4**).

**Figure 3.**
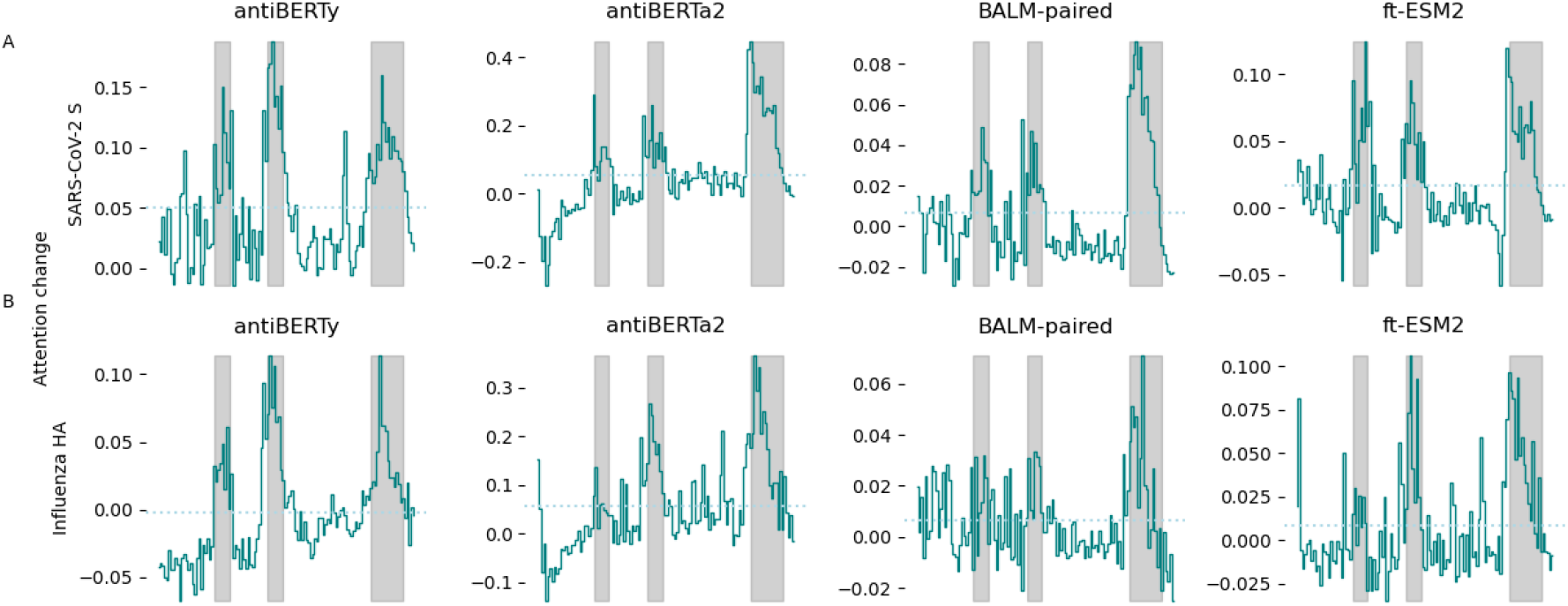
Change in average intra-heavy chain attention after fine-tuning. Attention activations were extracted from pre-trained and fine-tuned language models for 50 randomly selected antibodies specific for SARS-CoV-2 S protein **(A)** and influenza HA **(B)**, respectively. The intra-heavy chain attention activations were averaged across heads and layers for each position on the heavy chain. Differences in average attention activations before and after finetuning for the last three layers were computed. The x-axis represents the position along the heavy chain. The solid line indicates the mean change in average attention activations across the 50 antibodies. The gray background indicates the regions spanned by CDR of the antibodies. The dotted line represents the mean of the difference in attention activations.

### Fine-tuned specificity classifiers capture changes in repertoire following vaccination

To further evaluate if the fine-tuned language model classifiers capture specificity information, we applied the classifiers to two single-cell BCR repertoire datasets measuring immune response to SARS-CoV-2 vaccination and influenza vaccination [16,18].

In the SARS-CoV-2 mRNA vaccination dataset, eight donors had samples taken from two different tissues at various time points after SARS-CoV-2 vaccination: peripheral blood plasmablasts taken one week after the second immunization (Day 28) and axillary lymph node samples taken one to twelve weeks after vaccination (Day 28, Day 35, Day 60, Day 110). We first applied the fine-tuned language model S protein classifiers on individual sequences of the SARS-CoV-2 vaccination dataset and a control peripheral blood dataset, which were taken pre-pandemic and assumed to have low level of S protein-specific sequences, to see if the S protein classifiers can capture the immune response. We excluded any sequences within the same clone of the sequences in our training dataset to prevent data leakage and only kept one sequence from each clone to weigh each clone equally. The SARS-CoV-2 vaccination repertoires are similar in distribution of gene usage, CDR3 length and somatic hypermutation frequency with the control samples (**Figure S5**). We then averaged the predicted class probability from the S protein classifiers. We tested the difference in the mean predicted probability of binding to S protein between the peripheral blood plasmablast data and the control datasets using a Wilcoxon rank-sum test and found a significantly higher mean predicted probability of binding to S protein for the samples after SARS-CoV-2 vaccination repertoires (**Figure 4A)**, which matches with the plasmablast response after vaccination. Similarly, we applied the S protein classifiers to the lymph node repertoires after SARS-CoV-2 vaccination and computed the mean predicted probability. We found a persistent level of the mean predicted probability across the timepoints, which is also consistent with the robust and persistent germinal center response observed after two doses of the SARS-CoV-2 vaccination (**Figure 4B**).

**Figure 4.**
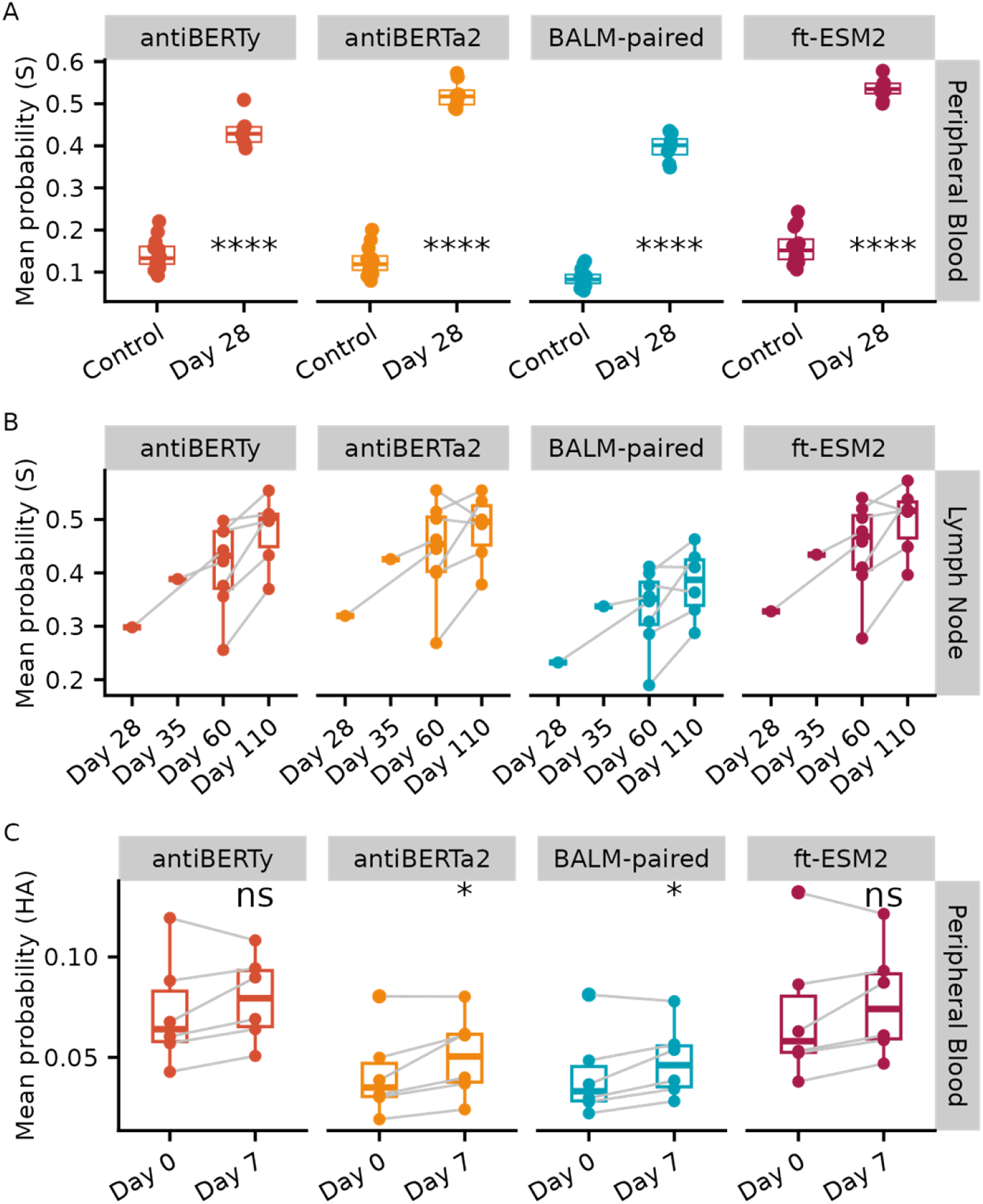
Application of the fine-tuned language model-based classifiers to vaccine response repertoire datasets. **(A)** Mean predicted probability of SARS-CoV-2 S protein binders by finetuned language model S protein classifier of the receptors from peripheral blood samples 28 days after SARS-CoV-2 vaccination (Day 28), compared with the pre-pandemic repertoire datasets (Control). Samples from the same donor were connected by lines. Paired Wilcoxon rank sum test was used to obtain the significance level of the increase in mean predicted probability at day 7 (ns: p > 0.05, *: p <= 0.05, **: p <= 0.01, ***: p <= 1e-3, ****: p <= 1e-4). **(B)** Mean predicted probability of S protein binders applied to lymph node repertoires 28, 35, 60, 110 days after SARS-CoV-2 vaccination. **(C)** Mean predicted probability of influenza HA binders by language model fine-tuned on HA classification task applied to peripheral blood repertoires before and seven days after influenza vaccination.

We used the same criterion to process the influenza vaccination repertoire datasets, which consisted of six influenza vaccine-responsive donors with peripheral blood samples taken at prevaccination (Day 0) and seven days post-vaccination (Day 7) for paired BCR heavy and light chain sequencing. We similarly applied the fine-tuned language model classifiers to individual sequences within the repertoires to compute predicted probability of individual sequences of binding to HA and average the predicted probability for each sample (**Figure 4C**). We performed Wilcoxon-rank sum tests between the two timepoints, and found an increase in average predicted class probability at Day 7 for all four fine-tuned models, with antiBERTa2 and BALM-pair significantly increased by paired Wilcoxon-rank sum tests, which is consistent with the observed antibody titer increases based on the HAI [16].

## Discussion

In this study, we investigated the efficacy of supervised fine-tuning on pre-trained antibody language models to improve specificity prediction to two key antigens, the SARS-CoV-2 S protein and influenza HA. We established a performance baseline using nested cross-validation of SVM models on pre-trained language model embeddings, showing improved performance with full-length input receptor sequences as opposed to just CDR regions. We compared the performance of fine-tuned models with supervised classifiers trained on embeddings from the same pre-trained language models and found that fine-tuning the language models led to enhanced specificity prediction. Additionally, we observed increased attention at the CDR regions after fine-tuning, indicative of the models capturing relevant features for antigen specificity. Furthermore, we applied the fine-tuned classifiers to longitudinal paired BCR repertoire data related to influenza and SARS-CoV-2 vaccination, showing their ability to capture changes in repertoire following vaccination, as evidenced by shifts in predicted binding probabilities.

While our study provides valuable insights into the effectiveness of fine-tuning pre-trained antibody language models for antigen specificity prediction, several limitations should be acknowledged. Firstly, the performance of the fine-tuned models may be influenced by the size and composition of the training datasets, such as the low frequency of ground truth non-binding sequences which could affect the generalizability of our findings to different datasets or tasks [19]. Secondly, our evaluation focused primarily on two specific antigens, the SARS-CoV-2 S protein and influenza HA, limiting the broader applicability of our conclusions to other antigens or biological contexts. Our study also did not explore the use of information on different epitopes of antigens, leaving room for future investigations to explore these avenues for improvement by leverage approaches such as multi-task learning. Additionally, the interpretation of attention activations changes after fine-tuning may be constrained by the complexity of attention mechanisms in language models since this extracted pattern may not have a straightforward mapping with the interactions between amino acid residues. Future studies on ground truth data are needed to further examine the utility of attention patterns for interpretability. More generalizable methods, including linguistics-inspired experimentation and grammatical inference, has been suggested as potential approaches to extracting sequence-function rules that the model has learned [20].

In summary, our study demonstrates the efficacy of fine-tuning pre-trained antibody language models to enhance specificity prediction. We established performance baselines and observed improved prediction accuracy with fine-tuned models, particularly in capturing changes in repertoire following vaccination. The findings give insights for further studies to advancing our understanding of antigen specificity prediction applications using antibody language models.

## Materials and Methods

### Models

We downloaded the following four pre-trained antibody language models, with the size parameters listed in **Table 1**.

**antiBERTy** [1]: Pre-trained using the BERT architecture on 588 million unpaired antibody heavy and light chain sequences from multiple species using a masked language modeling (MLM) objective.

**antiBERTa2** [3]: Based on the RoFormer architecture, pre-trained with 1.54 billion unpaired and 2.9 million paired human antibody sequences with MLM objective.

**BALM-paired** [6]: Developed using a RoBERTa-large architecture trained on 1.34 million paired antibody sequences with MLM objective.

**ft-ESM2** [6]: Based on 650-million parameter ESM2 (Evolutionary Scale Modeling) model [11], fine-tuned with 1.34 million paired antibody sequences with MLM objective.

### Data sources

We collected antibody sequences with specificity labels to influenza HA protein and SARS-CoV-2 S protein from public sources, as listed in **Table 2**.

#### Influenza HA-specific sequences

We extracted the paired-chain antibody sequences with influenza HA proteins binding/non-binding labels from public datasets [18,21,22], which consisted of 3,221 sequences binding to various HA proteins as well as 706 were non-binding. To balance the labels, we sampled additional vaccine non-responsive sequences from six pre-vaccination repertoires in [16] as additional negative controls. The distribution in V, J gene usage and CDR3 length is similar between the negative controls and the positive sequences (**Figure S1**). In total, 6,424 receptors were available, with 3,221 binding (50.1 %).

#### SARS-CoV2 S protein-specific sequences

We used the antibody sequences dataset we previously curated with binding labels to SARS-CoV-2 [15].

#### Repertoire data

We collected additional single-cell paired-chain repertoire data from [16], which had peripheral blood samples collected before and seven days after influenza vaccination, as well as [18], which had both peripheral blood and lymph node samples taken from 28, 35, 60 and 110 days after SARS-CoV-2 vaccination.

### Receptor specificity prediction using pre-trained language model embedding

To establish a baseline performance for the four language models in predicting specificity to the SARS-CoV-2 S protein and influenza HA proteins, we trained supervised models using the pre-trained model embedding as input. The process involved concatenating each pair of BCR heavy and light chain sequences, separated by two [CLS] tokens, and feeding them into each pre-trained antibody language model to obtain the output from the last hidden layer. Then, utilizing this embedding as input, we trained separate support vector machine (SVM) classifiers to predict the binary binding status for each antigen from each pre-trained model embedding.

Specifically, we employed sklearn SVM with an RBF kernel and implemented nested cross-validation to split the data into training, validation, and test sets, ensuring non-overlapping donors and preserving class percentage with sklearn.model_selection.StratifiedGroupKFold. Three inner loops and four outer loops were utilized for hyperparameter search on the validation set and to compute test set performance, respectively. During hyperparameter search, we conducted a grid search over the regularization parameter C of SVM, ranging from 0.01 to 100, and selected the optimum value based on the validation set AUROC score.

Evaluation of the test set performance included metrics such as AUROC, weighted-average F1 score, precision, recall, average precision score, balanced accuracy, and Matthews correlation coefficient. Finally, we chose the regularization parameters that yielded the best validation AUROC across nested CV outer folds and trained the final classifier using all available binding data.

### Supervised Fine-tuning of language models for receptor specificity prediction

We fine-tuned the last three layers of each of the four language models to predict the binary binding or non-binding status to either the SARS-CoV-2 S protein or influenza HA proteins. We assessed the performance of this fine-tuning by using the same cross-validation train-test data split employed in the embedding SVM approach for direct comparison. For each training dataset, we separated out a validation set (33%) to determine the optimum epoch. To fine-tune each language model, we instantiated a sequence classification model using transformers.AutoModelForSequenceClassification) and initialized it with the pre-trained weights for each model in **Table 1**. We trained each classification model with a learning rate of 1e-5, a batch size of 64 for 30 epochs, and selected models from epochs with the best validation AUROC to evaluate the test set performance by AUROC, weighted-average F1 score, precision, recall, average precision score, balanced accuracy, and Matthews correlation coefficient. We determined the epoch that yielded the best average validation AUROC across outer folds and trained the final classifier using all available binding data. All models were fine-tuned on a single NVIDIA RTX A5000 GPU.

### Applying sequence specificity classifiers to repertoires

To determine whether the classifier effectively identifies BCR specificity, we applied the classifiers to paired-chain BCR repertoire data from vaccinations against SARS-CoV-2 and influenza. In processing these datasets, we used the immcantation Change-O pipeline [24] to cluster BCR sequences into clonal groups. To prevent data leakage, we excluded sequences from the repertoires that belonged to the same clone as those used in training the specificity classifiers. Additionally, to minimize the confounding effects of clonal expansion, we retained only one sequence from each clone. For each sequence, we calculated the predicted class probability of binding to a given antigen and then computed the average of these predicted probabilities for each repertoire.

## Data and code availability

All data were from public sources as listed in **Table 2**. We deposited the code at https://bitbucket.org/kleinstein/projects/src/master/Wang2024/. Both code and data were deposited on Figshare at https://doi.org/10.6084/m9.figshare.25342924.

## Acknowledgements

We thank Katherine McIntire and Ali Ellebedy for sharing data, Hailong Meng for data processing, and Gisela Gabernet for manuscript discussion.

